# The Impact of Psychopathology, Social Adversity and Stress-relevant DNAm on Prospective Risk for Post-traumatic Stress: A Machine Learning Approach

**DOI:** 10.1101/2020.09.25.313635

**Authors:** Agaz H. Wani, Allison E. Aiello, Grace S. Kim, Fei Xue, Chantel L. Martin, Andrew Ratanatharathorn, Annie Qu, Karestan Koenen, Sandro Galea, Derek E. Wildman, Monica Uddin

**Author notes:** Corresponding Author: Monica Uddin, Ph.D. Genomics Program, College of Public Health University of South Florida, Tampa, FL 33612.

## Abstract

**Background:** A range of factors have been identified that contribute to greater incidence, severity, and prolonged course of post-traumatic stress disorder (PTSD), including: comorbid and/or prior psychopathology; social adversity such as low socioeconomic position, perceived discrimination, and isolation; and biological factors such as genomic variation at glucocorticoid receptor regulatory network (GRRN) genes. This complex etiology and clinical course make identification of people at higher risk of PTSD challenging. Here we leverage machine learning (ML) approaches to identify a core set of factors that may together predispose persons to PTSD.

**Methods:** We used multiple ML approaches to assess the relationship among DNA methylation (DNAm) at GRRN genes, prior psychopathology, social adversity, and prospective risk for PTS severity (PTSS).

**Results:** ML models predicted prospective risk of PTSS with high accuracy. The Gradient Boost approach was the top-performing model with mean absolute error of 0.135, mean square error of 0.047, root mean square error of 0.217, and R^2^ of 95.29%. Prior PTSS ranked highest in predicting the prospective risk of PTSS, accounting for >88% of the prediction. The top ranked GRRN CpG site was cg05616442, in *AKT1*, and the top ranked social adversity feature was loneliness.

**Conclusion:** Multiple factors including prior PTSS, social adversity, and DNAm play a role in predicting prospective risk of PTSS. ML models identified factors accounting for increased PTSS risk with high accuracy, which may help to target risk factors that reduce the likelihood or course of PTSD, potentially pointing to approaches that can lead to early intervention.

## Introduction

Post-traumatic stress disorder (PTSD) is a common and severe psychiatric disorder that develops following exposure to life-threatening or terrifying events (Bisson, Cosgrove, Lewis, & Robert, 2015; Shalev, 2001). It can develop following exposure to a single horrifying event or a prolonged exposure to a series of traumatic events such as physical or sexual assault, or combat. Not all people develop PTSD following exposure to trauma. Many people show the ability to recover from trauma exposure (Bonanno, 2004). Nevertheless, in a subset of individuals, PTSD can be severe and debilitating and show a chronic course over time (Perkonigg et al., 2005; Zvolensky et al., 2015). In addition, individuals with PTSD show different symptom presentations (Galatzer-Levy & Bryant, 2013) and often exhibit comorbid psychopathology, including depression and anxiety (Ginzburg, Ein-Dor, & Solomon, 2010). Comorbid psychopathology is likely to affect recovery and treatment outcome (Bradley, Greene, Russ, Dutra, & Westen, 2005; Dalenberg, Glaser, & Alhassoon, 2012). Importantly, despite many decades of research, it remains challenging to predict individuals at high risk of prospective post-traumatic psychopathology.

It is well-established that social adversity, such as low socioeconomic position (SEP), isolation, and discrimination, have a significant impact on mental and physical health. Multiple studies have shown a strong association between SEP and increased risk of psychopathology (Koenen, Moffitt, Poulton, Martin, & Caspi, 2007; Uddin et al., 2011; Ward-Caviness et al., 2020). People with low SEP are at a higher risk of mental health problems (Kiely, Leach, Olesen, & Butterworth, 2015; Ramos-Lima, Souza, Teche, & Freitas, 2019) and low SEP can adversely impact access to treatment and prevention of mental health conditions. In addition to SEP, research shows that parental education, and gender are associated with a prospective or higher risk of psychopathology (Park, Fuhrer, & Quesnel-Vallée, 2013). Similarly, social relationships or isolation/loneliness profoundly affect mental and physical health, including risk for PTSD (Cacioppo, Cacioppo, Capitanio, & Cole, 2015; Hyland et al., 2019; Kuwert, Knaevelsrud, & Pietrzak, 2014; Link & Phelan, 1995). Also, the contribution of perceived discrimination, which can engender feelings of isolation, has been significantly associated with many health conditions, including PTSD (Brooks Holliday et al., 2018; Kessler, Mickelson, & Williams, 1999), with people experiencing perceived discrimination more likely to show symptoms of the disorder (Bogart et al., 2011).

PTSD is thus shaped by a complex mix of stressful and traumatic life events that shape how our body responds to stress. Our central stress response system, the hypothalamic pituitary adrenal (HPA) axis plays a key role here. The HPA axis involves a complex set of instructions and feedback interactions between the hypothalamus, pituitary, and adrenal gland. Its main job is distributing glucocorticoid hormones, such as cortisol released by the adrenal gland (Whitnall, 1993). These hormones are vital for life and play a key role in mediating the stress response of the HPA axis. Numerous studies have shown that dysregulation in HPA axis function involving glucocorticoids, e.g., cortisol, plays a crucial role in PTSD pathophysiology (Rachel Yehuda, Halligan, Golier, Grossman, & Bierer, 2004; R. Yehuda et al., 1993). Dysregulation in the HPA axis has also been associated with major depressive disorder (Anacker, Zunszain, Carvalho, & Pariante, 2011; Menke et al., 2012; Rachel Yehuda et al., 2004). Genomic variation at glucocorticoid receptor regulatory network (GRRN) genes has been associated with PTSD (Binder et al., 2008; Labonté, Azoulay, Yerko, Turecki, & Brunet, 2014), childhood maltreatment (Bustamante et al., 2016), and depression (Bustamante et al., 2016; Hodes, Ménard, & Russo, 2016). These studies demonstrate that stress-relevant genomic and epigenomic variation, including DNAm variation, may play a role in predicting stress-related psychopathology.

As previously noted, predicting prospective risk of psychopathology remains a challenging task. Recent work, however, has seen a burgeoning interest in the application of machine learning (ML) methods for predicting PTSD risk (Karstoft, Galatzer-Levy, Statnikov, Li, & Shalev, 2015; Wshah, Skalka, & Price, 2019). ML refers to those approaches that do not use explicit programming but instead learn from data and experience to make decisions or predictions. For example, ML has recently been used to differentiate between combat-related PTSD and trauma-exposed controls (Zhang, Richardson, & Dunkley, 2020), and for diagnosing PTSD (Dean et al., 2019). With respect to PTSD, identifying factors that prospectively predict PTSD can potentially help target such factors to reduce the likelihood of PTSD following a traumatic event, and/or reduce the likelihood of a more chronic course of PTSD. To date, however, existing work has not considered social adversity-related factors, including loneliness and perceived discrimination, in developing ML-based models predicting PTSD, or combined such data with DNAm measures of relevance to stress-related psychopathology. To address these gaps, here we build ML models that incorporate factors previously associated with elevated risk of PTSD, including prior psychopathology, social adversity exposure, and DNAm variation in GRRN genes to predict prospective risk of PTSD symptom severity (PTSS).

## Materials and Methods

In this study, we used data from the Detroit Neighborhood Health Study (DNHS), a prospective population-based longitudinal cohort of individuals living in Detroit, Michigan (Goldmann et al., 2011; Uddin et al., 2010). All participants in this study were 18 years or older and predominantly self-identified as African American (AA). The main aim of the DNHS was to identify how genetic variation, stressful and traumatic life experiences, and features of the environment predict psychopathology and behavior. Participants were recruited for a structured telephone interview each year between 2008-2013 to assess perceptions of participant’s neighborhoods, mental and physical health status, social support, exposure to traumatic events, posttraumatic stress disorder symptoms, depression symptoms, generalized anxiety symptoms and alcohol, and tobacco use. Informed consent was obtained at the beginning of each interview and again at specimen collection. The Institutional Review Board of the University of Michigan and the University of North Carolina-Chapel Hill reviewed and approved this study.

### Measures

Demographic information, such as age, sex, race, education, marital status, and employment was self-reported by the participants. PTSD was assessed according to DSM-IV criteria using the PTSD Checklist Civilian Version (PCL-C) as described in (Uddin et al., 2010). Exposure to lifetime traumatic events was assessed using a survey of 19 item traumatic events, as in previous work (Breslau et al. 1998). Cumulative traumatic burden was estimated by summing the scores of lifetime traumatic event types (Uddin et al., 2010). Similarly, major depressive disorder (MDD) and generalized anxiety disorder (GAD) were measured using Patient Health Questionnaire (PHQ-9) and GAD-7 scales, respectively, as described earlier (Uddin et al., 2010). In addition to the categorical diagnosis of PTSD, we used continuous measures of PTSD, based on summing scores for 17 symptoms from the worst lifetime trauma. Also, severity measures for PTSD symptom clusters (intrusion, avoidance and hyperarousal) were used, by summing symptoms related to each symptom cluster. Depression and anxiety symptom severity measures based on respective scales (PHQ-9 and GAD-7) were used as well. Perceived discrimination was measured using the Everyday Discrimination Scale (EDS), a nine-item self-report scale (Williams, Yan, Jackson, & Anderson, 1997). Loneliness was measured using a three-item scale (Hughes, Waite, Hawkley, & Cacioppo, 2004). Emotional mistreatment, financial problems, legal issues, drug and alcohol-related problems, job loss, unemployment, divorce were measured as standalone stressors.

### DNAm collection, quality control, and pre-processing

Biospecimens were collected from DNHS participants who consented to give a sample. In this study, DNAm data were collected from DNHS participants exposed to one or more traumas, via DNA isolated from venipuncture blood draws as described in (Uddin et al., 2010).

DNAm derived was measured using Illumina’s Infinium MethylationEPIC BeadChip in 500 samples from 190 unique participants following the manufacturer’s recommended protocol. Metadata from trauma-exposed participants was randomized to minimize plate and chip mediated batch effects (Harper, Peters, & Gamble, 2013). Resulting DNAm data was then subjected to quality control (QC): Sex and genotype checks were performed to remove sex discordant samples using the *minfi* and *ewastools* R packages (Aryee et al., 2014; Heiss & Just, 2018). Raw DNAm beta values were obtained using the *minfi* R package (Aryee et al., 2014). QC was performed to filter poorly performing samples and probes. Samples with low signal intensity (i.e. mean signal intensity <2000 arbitrary units), or <50% of the overall median were also removed (Barfield, Kilaru, Smith, & Conneely, 2012) as were samples and probes with >10% missing values. Probes with detection p-value >0.01 were set to missing. Cross-reactive and polymorphic probes were removed (McCartney et al., 2016).

QC removed a total of 52 samples, leaving 448 samples from 179 participants (from four time points/waves) for subsequent analysis. For ML, we used samples from waves 1 and 2 to predict PTSS from wave 4, and included survey data from waves 1, 2, and 3 for model building (described below). From the set of 448 samples that passed QC, a total of 210 samples from 148 unique participants met the criteria for inclusion in ML analyses (Figure 1). Normalization of the methylation data was performed using the *Noob* approach implemented in the *minfi* R package (Aryee et al., 2014; Triche, Weisenberger, Van Den Berg, Laird, & Siegmund, 2013). *ComBat* adjustment was performed to reduce the likelihood of bias due to known batch effects using an empirical Bayesian framework implemented in *SVA* R package (Johnson, Li, & Rabinovic, 2006; Leek, Johnson, Parker, Jaffe, & Storey, 2012). Cell estimations were computed using the *IDOL* algorithm (Salas et al., 2018).

**Figure 1:**
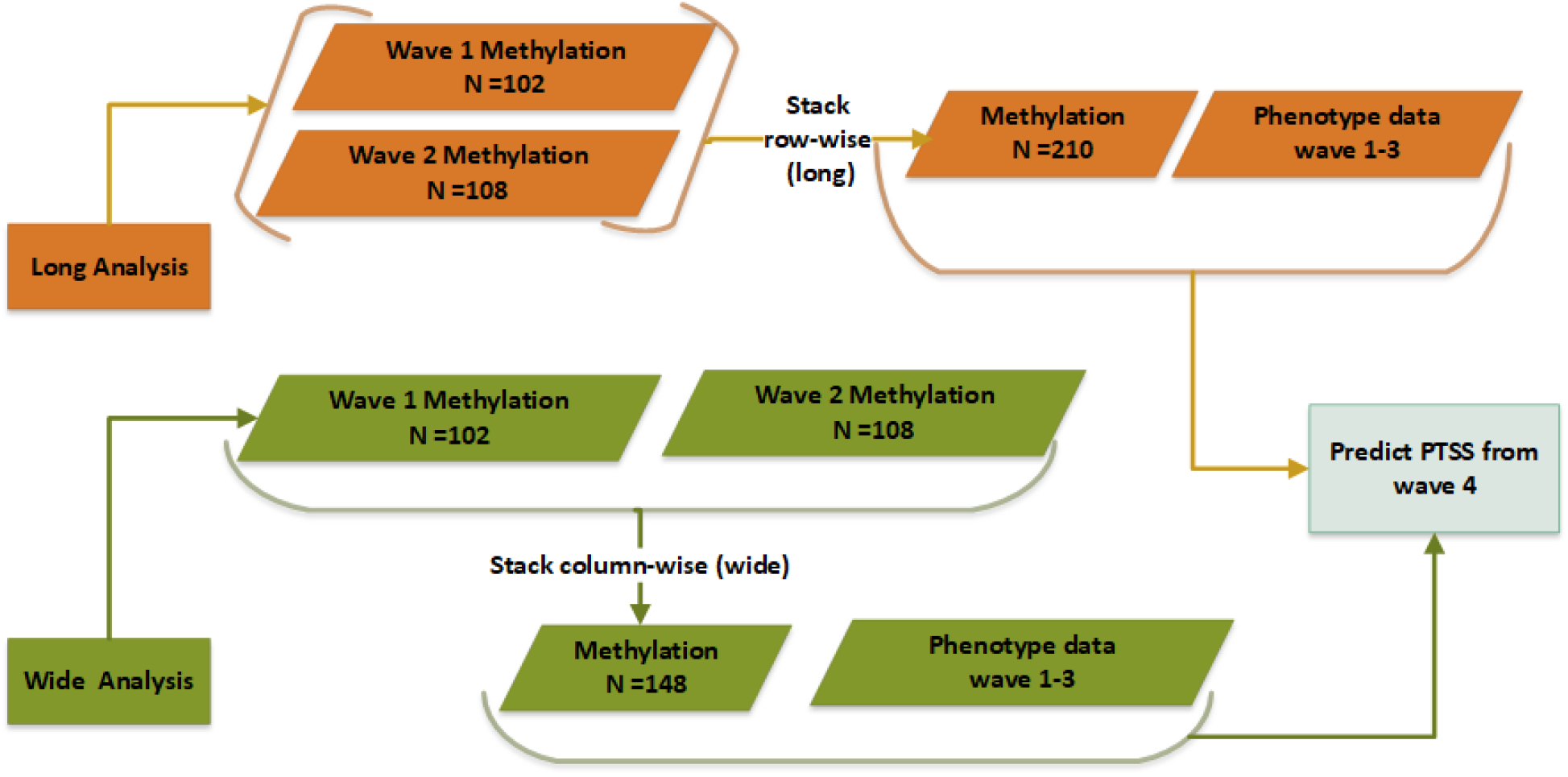
Two sets of analyses showing methylation data in a long and wide format, including the phenotype information. Long Analyses: combines methylation data from two-time points (waves) in a row-wise format (n = 210). Wide Analyses: combine methylation data from two-time points in a column-wise format (n = 148).

### Pre-processing for ML

To prepare the data (DNAm, phenotype and cell proportions) for ML, we performed some additional pre-processing steps, including imputation of missing data, feature selection, and scaling. The overall workflow of the pre-processing and ML models is shown in Figure 2.

**Figure 2:**
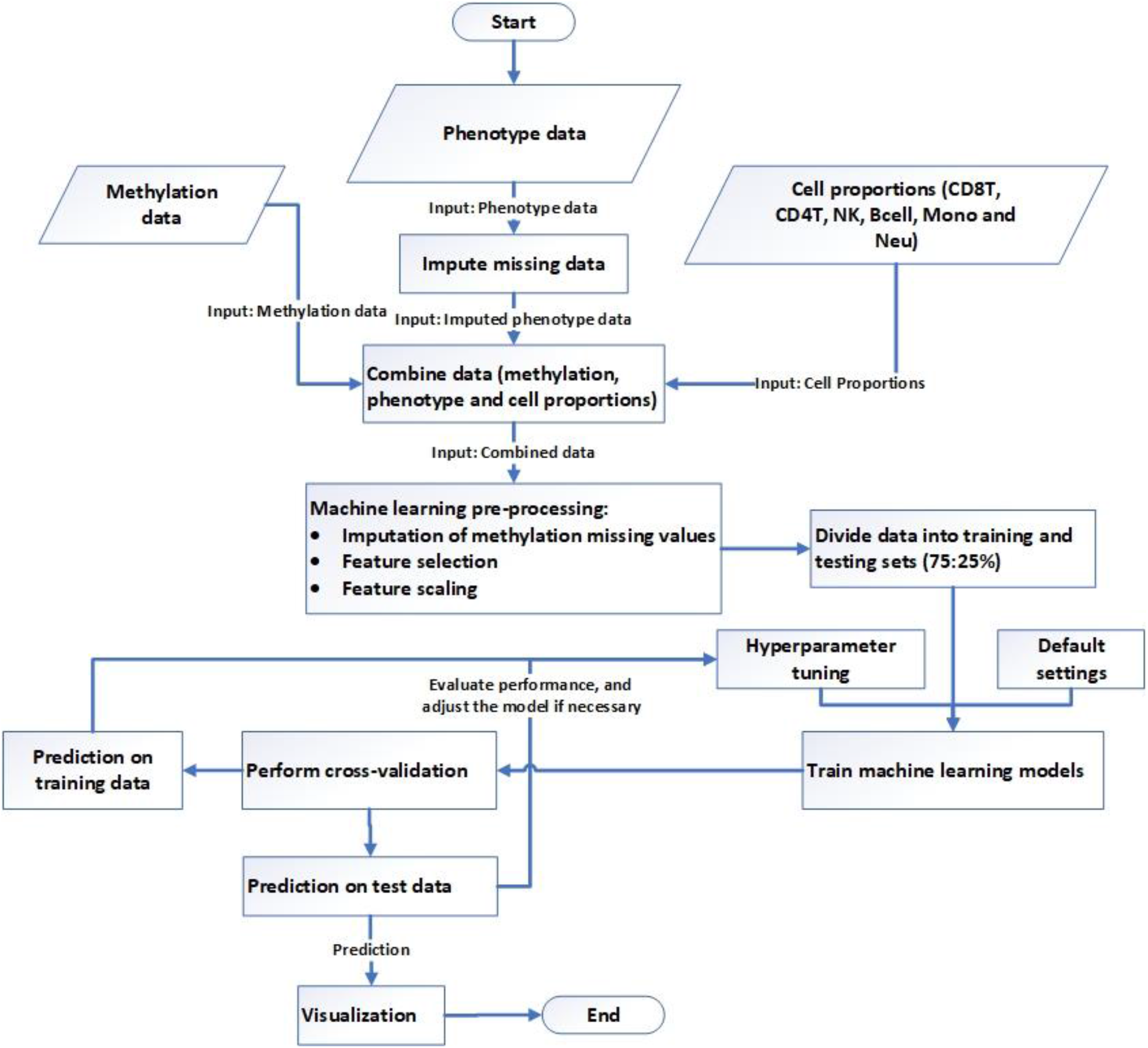
Workflow of the ML process. It starts with combining data (DNAm, phenotype and cell proportions) and ends with the PTSS risk prediction and visualization of the results.

#### General ML Approach

In this study, we used multiple well-known and efficient ML algorithms for prediction. As described in more detail below, the goal is to compare and determine the best performing algorithms predicting PTSD with high accuracy.

#### Imputation

As ML models require no missing data, we imputed both phenotype and DNAm data. We used Predictive Mean Matching (PMM), a semi-parametric approach to impute the phenotype information using the *mice* R package (van Buuren & Groothuis-Oudshoorn, 2011). DNAm data were imputed using the *k*-nearest neighbor approach (*k*=2) implemented in the *Scikit-learn* Python framework (Pedregosa et al., 2011; Troyanskaya et al., 2001). All the data (DNAm, cell estimation, and phenotypes) were combined for ML.

#### Feature selection

Feature selection is a critical pre-processing step used to remove redundant and irrelevant features; CpGs, phenotypes (including social adversity exposures) and estimated cell type proportions are all defined as features. This step helps to identify the significant features that are highly predictive for the outcome variable of interest. We used a univariate feature selection approach based on a univariate statistical test to select the top 150 features important for PTSS prediction, implemented in the *Scikit-learn* python framework (Pedregosa et al., 2011).

#### Feature Scaling

To standardize data, we performed feature scaling to standardize the data with mean 0 and standard deviation 1 using *Scikit-learn* framework (Pedregosa et al., 2011).

### Machine Learning Approaches

We used multiple approaches such as Random Forest (RF) (Breiman, 2001), Adaboost (AB) (Drucker, 1997; Freund & Schapire, 1997), Gradient Boost (GB) (Friedman, 2001), Linear Regression (LR) (Lai, Robbins, & Wei, 1978), Support Vector Regression (SVR) (Chang & Lin, 2011; Vapnik, 1995), Bagging Regression (BR) (Breiman, 1996) and Voting regression (VR) (An & Meng, 2010). More information about these approaches is given in Supplementary Information.

### Evaluation

#### Cross-validation

Cross-validation is a technique used to estimate how well a model will generalize to an independent test data set. We performed *k*-fold (*k*=10) cross-validation on the training data. It works by training the model on *k-1*-fold of the training data and validating on the *kth* fold. Each of the k-folds follows this approach, and average performance on *k*-folds is measured. Cross-validation is computationally intensive but does not require an independent validation dataset, thus making more data available for training.

To evaluate the error rate and accuracy of the models, we used standard error reporting metrics: mean absolute error (MAE), mean square error (MSE), root mean square error (RMSE), and R Squared (R^2^) (Supplementary Information). As we scaled the data, including the response variable, the error rates (MAE, MSE, and RMSE) are also scaled.

### Analysis

In the current sample from the DNHS, we have data from a baseline wave (wave 1) and three follow-up waves (wave 2, wave 3, and wave 4). We performed two types of analyses using ML. For both types of analysis, we used DNAm data from waves 1 and 2, and phenotype data from waves 1, 2, and 3 to predict PTSS from wave 4 as shown in Figure 1. In the first analysis, DNAm data from waves 1 and 2 were stacked together, such that participants with DNAm data from more than one wave were included as rows in the data matrix. This analysis included 210 samples from 148 unique participants. In the second analysis, we arranged participants with DNAm data from more than one wave as columns in the data matrix, to identify the CpGs and phenotypes that are significant in both waves. This analysis included 148 samples from 148 unique participants. In both analyses, we examined, using DNAm and phenotype data, how well we can predict the prospective risk of PTSS, and identify the CpG sites and phenotypes that are highly predictive for PTSS. From now onwards in this study, we will refer first analysis (DNAm row-wise) as “long” and the second as “wide” (DNAm column-wise) for the sake of simplicity.

As mentioned, feature selection is an important step in ML. To see if ML performs better on the data with a significant set of features as compared to the data with a full set of features, we used data in four different ways 1) All the DNAm features as long 2) All the DNAm features as wide 3) Important DNAm features as long and 4) Important DNAm features as wide. To implement the machine learning models, we divided the data into training and testing sets (75:25%).

## Results

We included a total of 210 samples from 148 unique DNHS participants for “long” ML analyses, and 148 samples from 148 unique participants for “wide” ML analyses. All participants were age 18 years or above and exposed to traumatic events. The participants predominantly self-identified as AAs (93.81%) and female (60%). The demographic characteristics of the cohort are shown in Table 1.

**Table 1:**
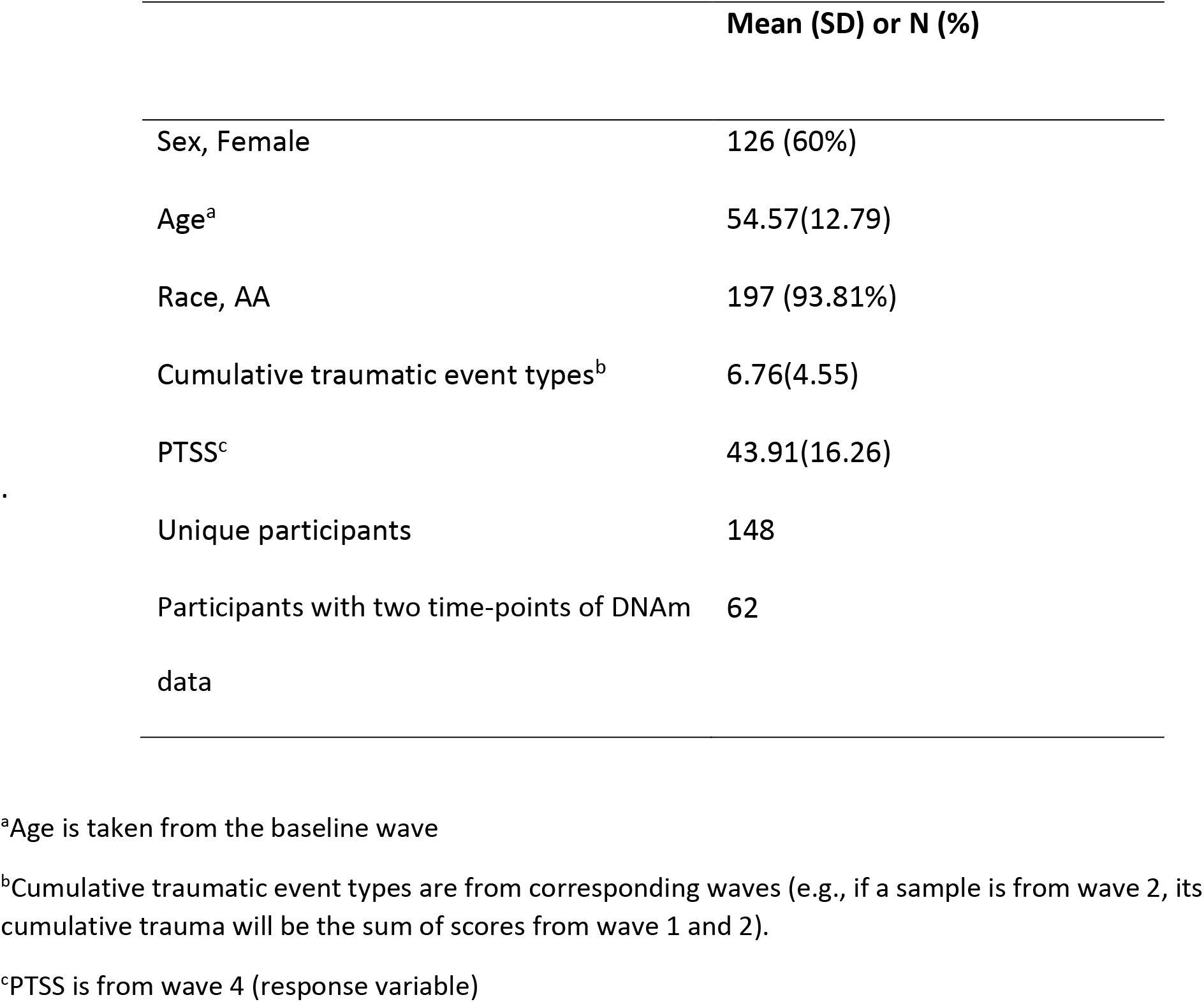
Demographic characteristics, traumatic events, PTSS score and number of unique participants in the study sample.

|  | Mean (SD) or N (%) |
| --- | --- |
| Sex, Female | 126 (60%) |
| Age<sup>a</sup> | 54.57(12.79) |
| Race, AA | 197 (93.81%) |
| Cumulative traumatic event types<sup>b</sup> | 6.76(4.55) |
| PTSS<sup>c</sup> | 43.91(16.26) |
| Unique participants | 148 |
| Participants with two time-points of DNAm data | 62 |

<sup>a</sup>Age is taken from the baseline wave
<sup>b</sup>Cumulative traumatic event types are from corresponding waves (e.g., if a sample is from wave 2, its cumulative trauma will be the sum of scores from wave 1 and 2).
<sup>c</sup>PTSS is from wave 4 (response variable)

### Best Features and Importance

First, to remove the irrelevant and redundant features, we performed feature selection and selected the top 150 features from both long and wide datasets, respectively. We performed a series of experiments to identify the set of features that most precisely predict PTSS. We tested sets of features that included between 10 - 590 features, increasing the size of feature set by increments of 20, until the error rate increased as we added more features. The models, RF, AB, and GB were used to find the errors (MAE and RMSE) of these different sized feature sets, and we evaluated the performance of these sets based on the mean of all three models. The feature set with the minimum mean error rate was used in the final analysis. Error rates of different feature sets are given in Table S1 in the supplementary file.

From the significant set of 150 features in the long data, 79 features were CpGs, and 71 were phenotype variables (Supplementary Figures S1 and S2). Similarly, from the wide data, 99 significant features were identified to be CpGs, while 51 were phenotype variables (Supplementary Figures S3 and S4). Between the long and wide data, 44 CpGs and 51 phenotypes were shared in common. Interestingly, three CpGs (cg04444450: *NCOA2*, cg20509117: *IL6; LOC541472* and cg05790989: *POU2F1)*, shown in Table 2, were consistently significant, meaning that they were in both waves of the wide dataset (wave 1 and wave 2) and in the long dataset. The important CpGs and phenotypes identified in both the long and wide datasets are shown in Figure 3. Prior PTSS ranked highest in predicting the prospective risk of PTSS, accounting for 88-89% of the prediction in the wide and long datasets, respectively. Psychopathology of depression and anxiety were also found to be significant in predicting PTSS risk. In addition to prior psychopathology, PTS symptom clusters (hyperarousal, intrusion, and avoidance) and social adversity factors (loneliness, perceived discrimination, financial problems, and emotional mistreatment) were significant predictors of PTSS risk. Using scores from both long and wide datasets, the top ranked GRRN CpG site was cg05616442 in the gene *AKT1* (the gene that encodes Protein kinase B, PKB) (see Table 2), and the top ranked social adversity feature was loneliness, followed by perceived discrimination and financial problems. The cumulative score of traumatic event types was also on the list of significant predictors for both datasets. In general, the importance of phenotypes ranked higher than the importance of most CpGs, although there were exceptions to this pattern (Figure 3; Figures S1-S4). Correlation of DNAm data in CpGs common to both approaches is shown in Figure 4. Most CpGs were positively correlated with each other; however, CpG cg26560981 in gene Mitogen-Activated Protein Kinase 10 (*MAPK10*) was negatively correlated with the greatest number of CpGs.

**Table 2:** Glucocorticoid receptor regulatory network CpGs and associated genes identified as important features in both the long and wide datasets*.

| **Illumina ID** | **UCSC_Ref Gene Name** | **Wide Importance** | **Long Importance** |
| --- | --- | --- | --- |
| cg05616442 | *AKT1* | 0.00418 | 0.00212 |
| cg18071894 | *AKT1* | 0.00281 | 0.00052 |
| cg26495008 | *FKBP5* | 0.00213 | 0.00037 |
| cg10913456 | *FKBP5* | 0.00196 | 0.00029 |
| cg23462257 | *NFKB1* | 0.00190 | 0.00199 |
| cg22237988 | *BAX* | 0.00175 | 0.00028 |
| cg26049684 | *AKT1* | 0.00153 | 0.00085 |
| cg25368824 | *IL4* | 0.00147 | 0.00356 |
| cg02842899 | *TP53;WRAP53* | 0.00144 | 0.00069 |
| cg02564102 | *SMARCC1* | 0.00137 | 0.00081 |
| cg11916669 | *AKT1* | 0.00127 | 0.00036 |
| cg13135255 | *CREBBP* | 0.00127 | 0.00054 |
| cg04444450 | *NCOA2* | 0.00110 | 0.00094 |
| cg24307117 | *SPI1* | 0.00097 | 0.00068 |
| cg23523922 | *AFP* | 0.00095 | 0.00028 |
| cg15912732 | *AKT1* | 0.00091 | 0.00074 |
| cg11540119 | *SMARCC1* | 0.00086 | 0.00040 |
| cg20509117 | *IL6;LOC541472* | 0.00084 | 0.00313 |
| cg24026230 | *NR3C1* | 0.00083 | 0.00057 |
| cg13986355 | *AKT1* | 0.00078 | 0.00065 |
| cg20730067 | *NFKB1* | 0.00074 | 0.00066 |
| cg14849556 | *NCOA1* | 0.00067 | 0.00146 |
| cg03883275 | *MAPK8* | 0.00057 | 0.00108 |
| cg19261497 | *MDM2* | 0.00057 | 0.00073 |
| cg16224829 | *NR3C1* | 0.00056 | 0.00081 |
| cg01277438 | *NFATC1* | 0.00047 | 0.00065 |
| cg11321922 | *NR1I3* | 0.00046 | 0.00014 |
| cg02521996 | *MAPK3* | 0.00039 | 0.00053 |
| cg23751680 | *SMARCA4* | 0.00038 | 0.00031 |
| cg13103915 | *CSF2* | 0.00038 | 0.00037 |
| cg26560981 | *MAPK10* | 0.00035 | 0.00068 |
| cg16569373 | *CREBBP* | 0.00033 | 0.00015 |
| cg09566021 | *CREBBP* | 0.00027 | 0.00194 |
| cg14825287 | *AKT1* | 0.00026 | 0.00134 |
| cg16182267 | *NFATC1* | 0.00026 | 0.00039 |
| cg20090430 | *GSK3B* | 0.00025 | 0.00041 |
| cg20509117 | *IL6;LOC541472* | 0.00023 | 0.00313 |
| **cg05790989** | *POU2F1* | 0.00019 | 0.00028 |
| **cg17406386** | *NR1I3* | 0.00017 | 0.00043 |
| **cg04444450** | *NCOA2* | 0.00011 | 0.00094 |
| **cg01312837** | *CREBBP* | 0.00010 | 0.00054 |
| **cg05121010** | *POU2F1* | 0.00010 | 0.00073 |
| **cg23624957** | *HDAC2* | 0.00008 | 0.00031 |
| **cg05790989** | *POU2F1* | 0.00004 | 0.00028 |

\*Shown are the Illumina ID for each CpG, UCSC gene name, and feature importance in the wide and long datasets.

**Figure 3:**
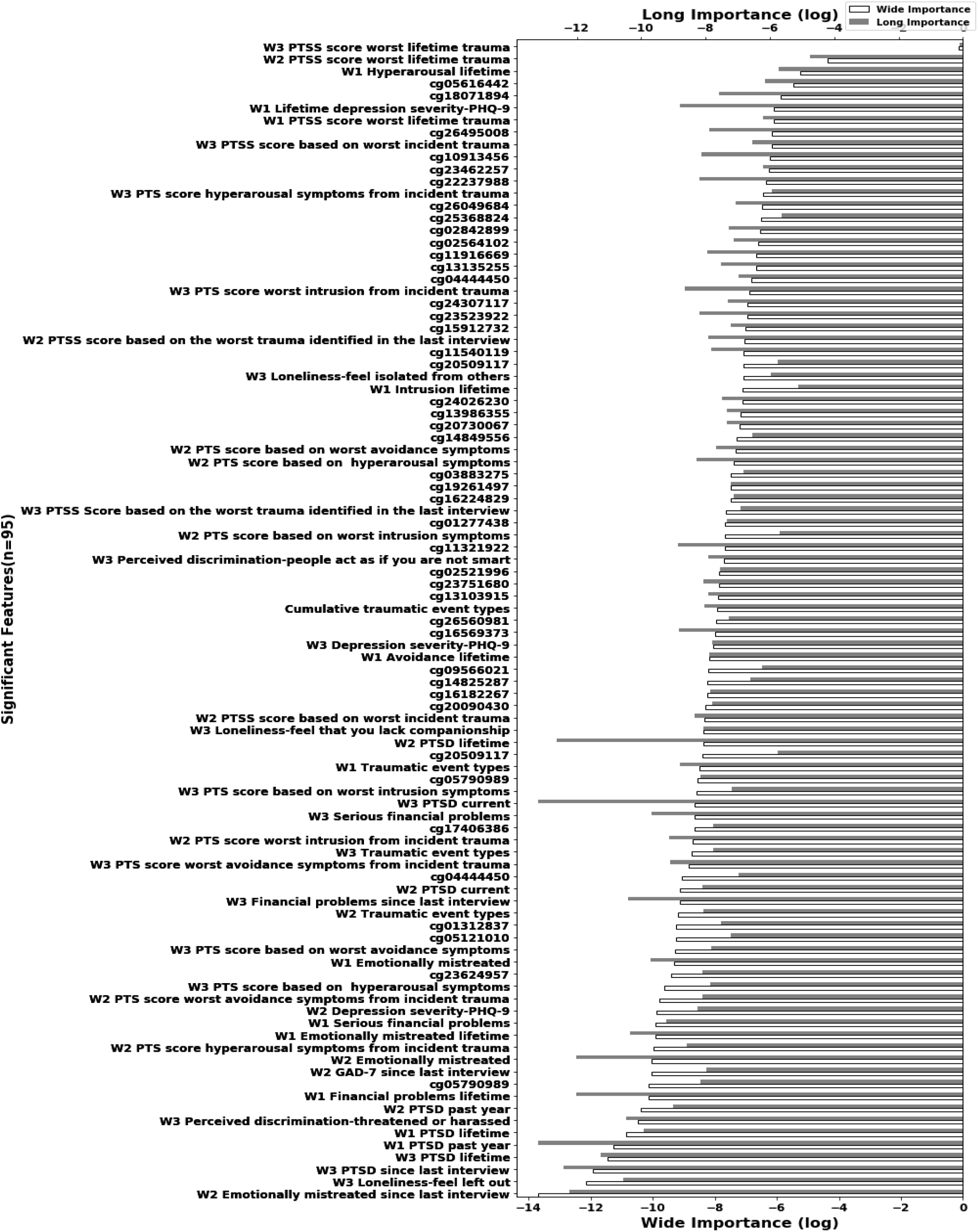
Features identified as important in both the long and wide datasets. Feature importance is in log scale, and the values near zero indicate higher importance. The importance of a significant feature can be between 0 and 1 and importance of all features sums to one. Filled bar: significance levels identified in the long dataset. White bar: significance levels identified in the wide dataset.

**Figure 4:**
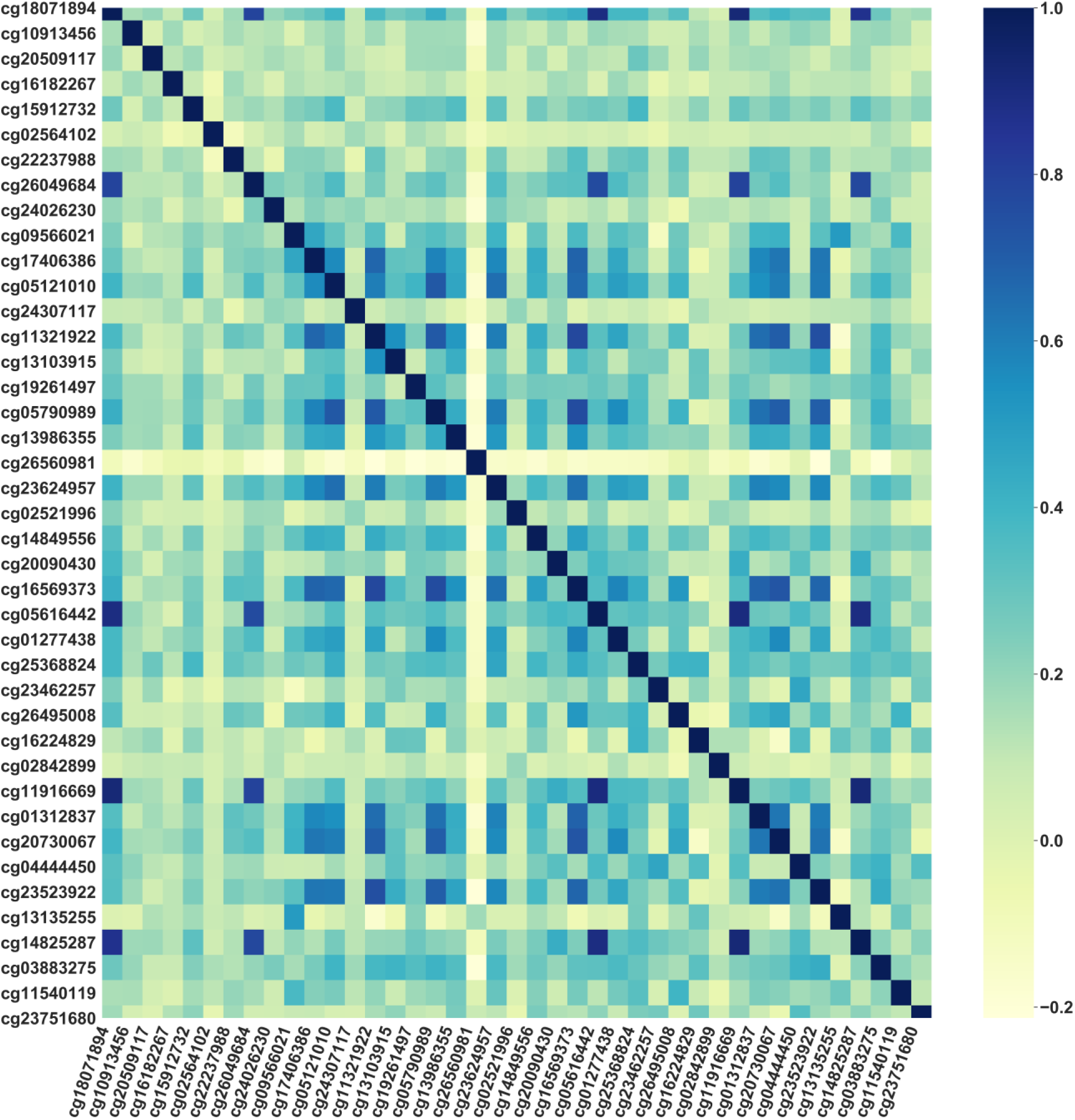
Correlation plot of DNAm values in CpGs identified as important in both long and wide analysis. Almost all the CpGs are positively correlated with each other. One CpG (cg26560981) shows a negative correlation with the largest number of CpGs.

### Prediction on Training Data and Cross-validation

Next, we used RF, AB, and GB to predict PTSS. These models were used in two different settings, i.e., the base model and tuned model, to search for the best set of parameters. In the base model, all the models used default settings provided by the *Scikit-learn* framework. After training the models in both the settings, we used 10-fold cross-validation on the training data to avoid overfitting of the models. Cross-validation gives us a sense of how well the model will generalize. The cross-validation error score of the base models on the training data is shown in Figure 5. For each model, we calculated the mean RMSE values. For both the long and wide datasets, the error score of the base models decreased when using just the set of significant features. The base models performed very well; cross-validation R^2^ values of the base models are shown in Figure 6. The mean R^2^ values for RF and AB were > 85%, and GB > 83%, which indicates the independent variables explain very well the variance of the dependent variable (PTSS). The prediction accuracies of all three models were close to each other; however, RF outperformed AB and GB. The mean RMSE on important features for RF = 0.331, AB = 0.344 and GB = 0.35 in long and RF = 0.344, AB = 0.359 and GB = 0.385 in wide data. Similarly, the R^2^ of the RF = 86.7% on long and 86.4% on wide, higher as compared to AB and GB. The cross-validation results using the base models were better on the significant set of features as compared to all the features, as shown in Figures 5 and 6.

**Figure 5:**
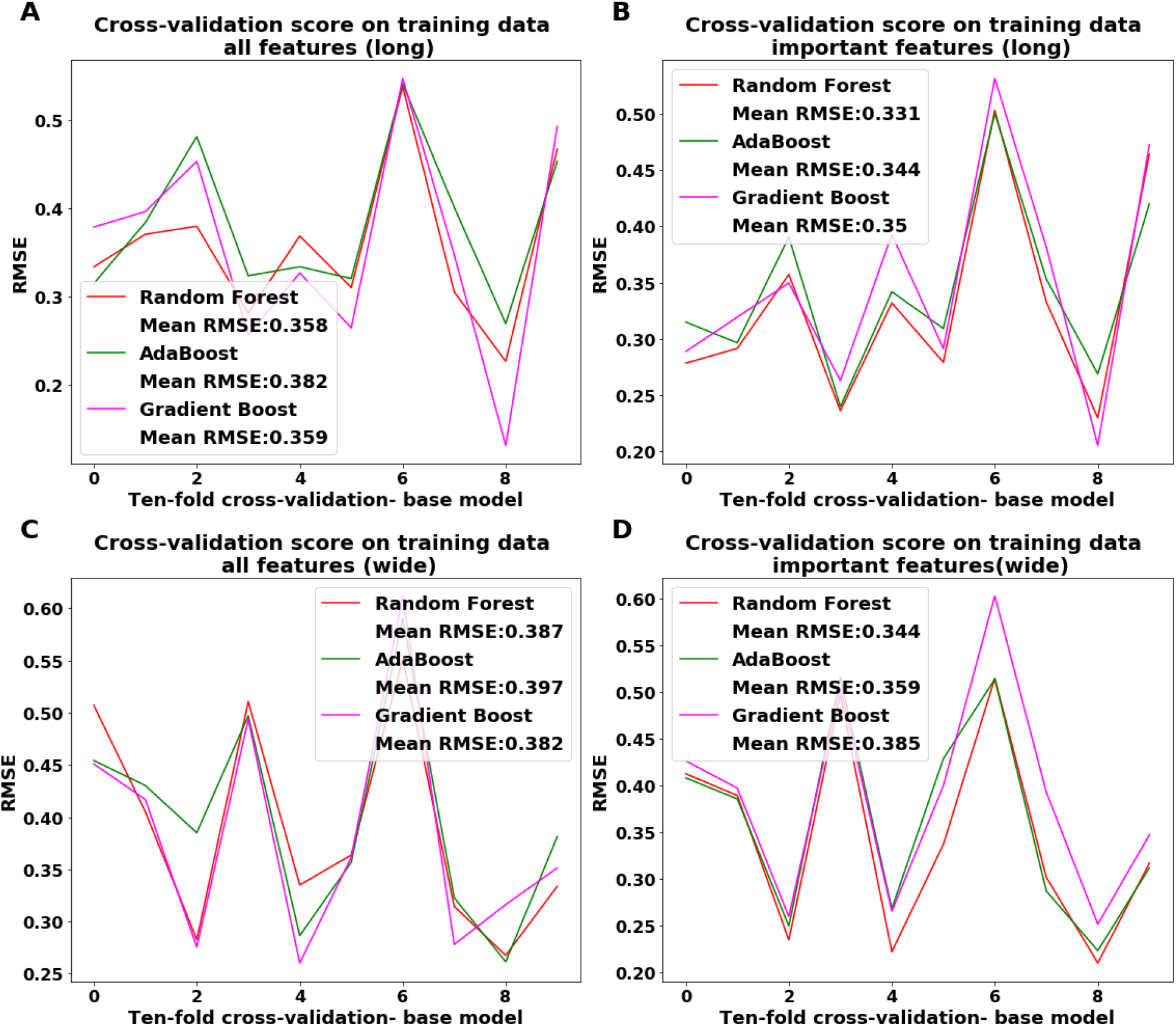
Ten-fold cross-validation on training data showing RMSE score on each of the ten-folds and mean RMSE on all ten-folds for each model. A) All features (GRRN genes, cell proportions and phenotype), and using DNAm features in long data. B) Important features (150) using DNAm in long. C) All features, using methylation in wide data. D) Important features, methylation in wide. The legend in each plot shows the mean RMSE on ten-fold cross-validation for each model. Using important features proved a better score (less RMSE) for both long and wide data.

**Figure 6:**
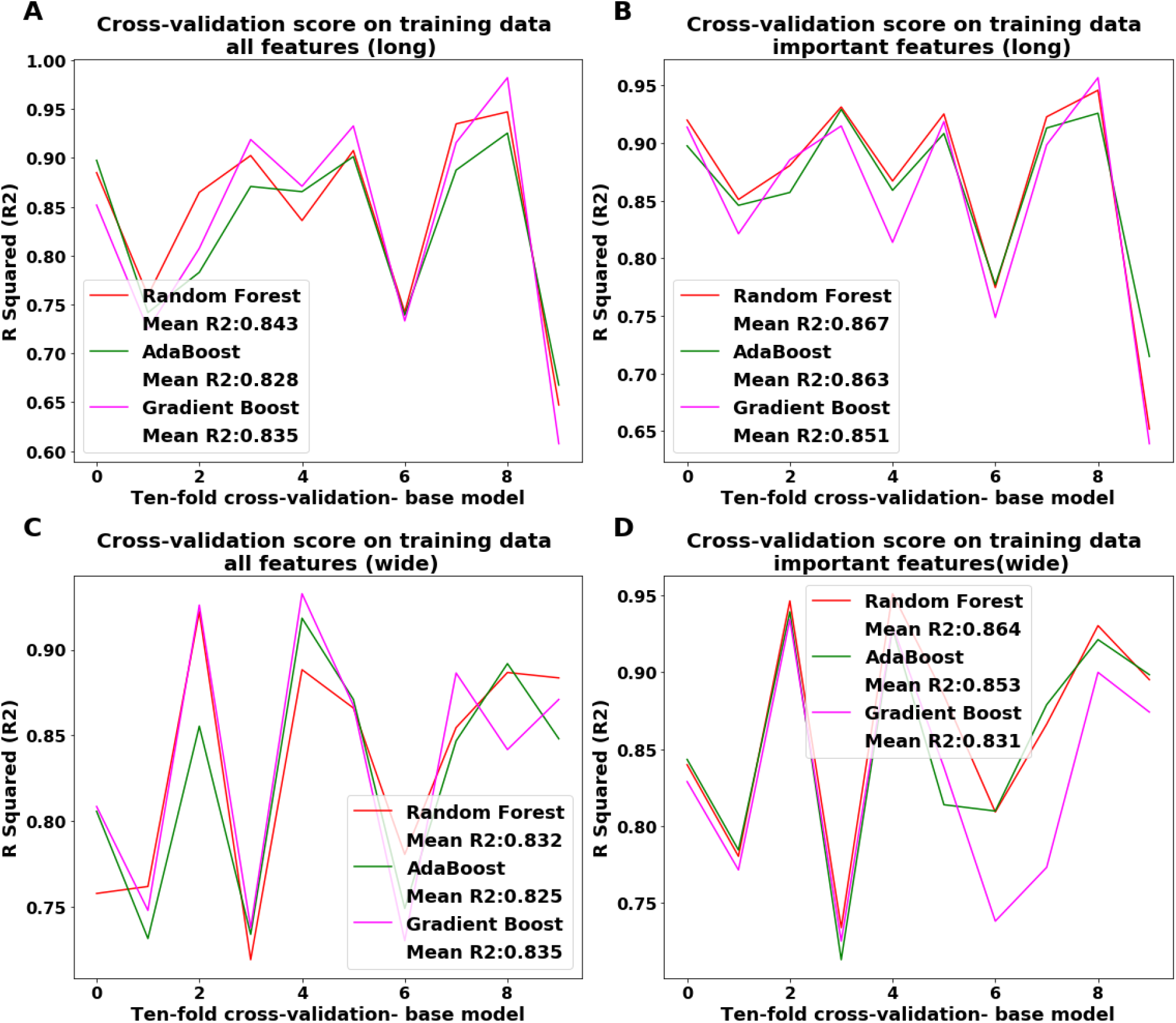
R squared (R^2^) values on each of the ten-folds and mean R^2^ for each model. The order of subplots A, B, C, D is the same as described in Figure 5. Results show better R^2^ value on an important set of features, both long and wide data, for all three models as compared to full data.

### Prediction Using Tuned Models

To potentially increase the performance of the base models, we ran 100 iterations to search for the best hyperparameters to tune the models. The results, RMSE and R^2^, are shown in the supplementary Figures S5 and S6, respectively. Tuning the hyperparameters did not increase the accuracy of the models. R^2^ for RF= 86.4% and GB= 85.1% in the long set; in fact, RF showed a small decrease in R^2^, suggesting the base model parameters are optimal. The AB showed a gain in R^2^ in the tuned model for both long (86.8%) and wide datasets (86%). Also, we recorded some gains for GB in the tuned model for the wide dataset. GB showed gain in tuned model accuracy, reducing the RMSE from .385 to .352 and increasing the R^2^ from 83.1% to 85%.

### Prediction on Test Data

After confirming the scores of base and tuned models using cross-validation to avoid over-fitting, we used the models on test data to get the final prediction score of the PTSS. The base model test score is shown in Table 3, and the tuned model score is shown in Table 4. It is interesting to note that all models provided a better score on the important long set of features in both the base and tuned models compared to the wide set of features There was a small difference when looking at the mean value of error rate and R^2^ of all three models in base (MAE: 0.2114, MSE: 0.0936, RMSE: 0.3025, R^2^: 0.9064) and tuned (MAE: 0.2056, MSE: 0.0916, RMSE: 0.3000, R^2^: 0.9083). The highest R^2^ values achieved on long test data using the base model and important set of features were GB = 95.29 %, followed by RF = 95.10%. Models performed better on the important set of features compared to all features in wide data.

**Table 3:** Performance measures on the test set using the base model.

| Model | Data | Mean Absolute Error | Mean Square Error | Root Mean Square Error | R Squared |
| --- | --- | --- | --- | --- | --- |
| Adaboost | Full long | 0.2128 | 0.0871 | 0.2952 | 0.9129 |
| Gradient Boost |  | **0.1830** | **0.0752** | **0.2742** | **0.9248** |
| Random Forest |  | 0.1779 | 0.0715 | 0.2674 | 0.9285 |
| Adaboost | Full wide | 0.2728 | 0.1420 | 0.3768 | 0.8580 |
| Gradient Boost |  | **0.2639** | **0.1204** | **0.3469** | **0.8796** |
| Random Forest |  | 0.2638 | 0.1248 | 0.3532 | 0.8752 |
| Adaboost | Important long | 0.2161 | 0.0834 | 0.2888 | 0.9166 |
| Gradient Boost |  | **0.1353** | **0.0471** | **0.2171** | **0.9529** |
| Random Forest |  | 0.1403 | 0.0490 | 0.2214 | 0.9510 |
| Adaboost | Important wide | 0.2452 | 0.1204 | 0.3470 | 0.8796 |
| Gradient Boost |  | **0.2059** | **0.0951** | **0.3084** | **0.9049** |
| Random Forest |  | 0.2205 | 0.1070 | 0.3271 | 0.8930 |

Full long: All the methylation features in the long data. Full wide: All methylation features in wide data. Important long: Important features in the long data. Important wide: important features in the wide data. All the three models performed best on the important set of features in long data. Results show GB to be the best performing approach.

**Table 4:** Performance measures on the test set using tuned model.

| Model | Data | Mean Absolute Error | Mean Square Error | Root Mean Square Error | R Squared |
| --- | --- | --- | --- | --- | --- |
| Adaboost | Full long | 0.1945 | 0.0778 | 0.2789 | 0.9222 |
| Gradient Boost |  | **0.1814** | **0.0742** | **0.2723** | **0.9258** |
| Random Forest |  | 0.1779 | 0.0727 | 0.2697 | 0.9273 |
| Adaboost | Full wide | 0.2518 | 0.1241 | 0.3523 | 0.8759 |
| Gradient Boost |  | **0.2282** | **0.0931** | **0.3051** | **0.9069** |
| Random Forest |  | 0.2605 | 0.1300 | 0.3605 | 0.8700 |
| Adaboost | Important long | 0.2111 | 0.0776 | 0.2786 | 0.9224 |
| Gradient Boost |  | 0.1672 | 0.0594 | 0.2437 | 0.9406 |
| Random Forest |  | **0.1473** | **0.0575** | **0.2399** | **0.9425** |
| Adaboost | Important wide | 0.2326 | 0.1088 | 0.3298 | 0.8912 |
| Gradient Boost |  | 0.1905 | 0.1156 | 0.3399 | 0.8844 |
| Random Forest |  | **0.2245** | **0.1088** | **0.3298** | **0.8912** |

Like the base model, tuned models performed best on the important set of features in long data. In tuned model RF approach performs best.

We also implemented many other models such as BR, LR, SVR, VR, the scores of which are shown in Table 5. As mentioned, the VR approach used all the models to get individual predictions and used the voting method for the final prediction. For all these models, we used the base model because hyperparameter tuning is computationally expensive and did not produce any notable gain (see Table 3 and Table 4). The bagging approach performed well and outperformed LR, SVR, and VR. All models performed well on an important set of features other than the LR approach, which performed poorly, see Table 5. When we compared all the methods on the test data, the base models on long and important features showed the best prediction as compared to the tuned models. The GB approach showed the best accuracy with MAE = 0.135, RMSE = 0.217, and R2 = 95.29%. The prediction of the RF approach was very close to GB with MAE = 0.140, RMSE = 0.221 and R2 = 95.10%.

**Table 5:**
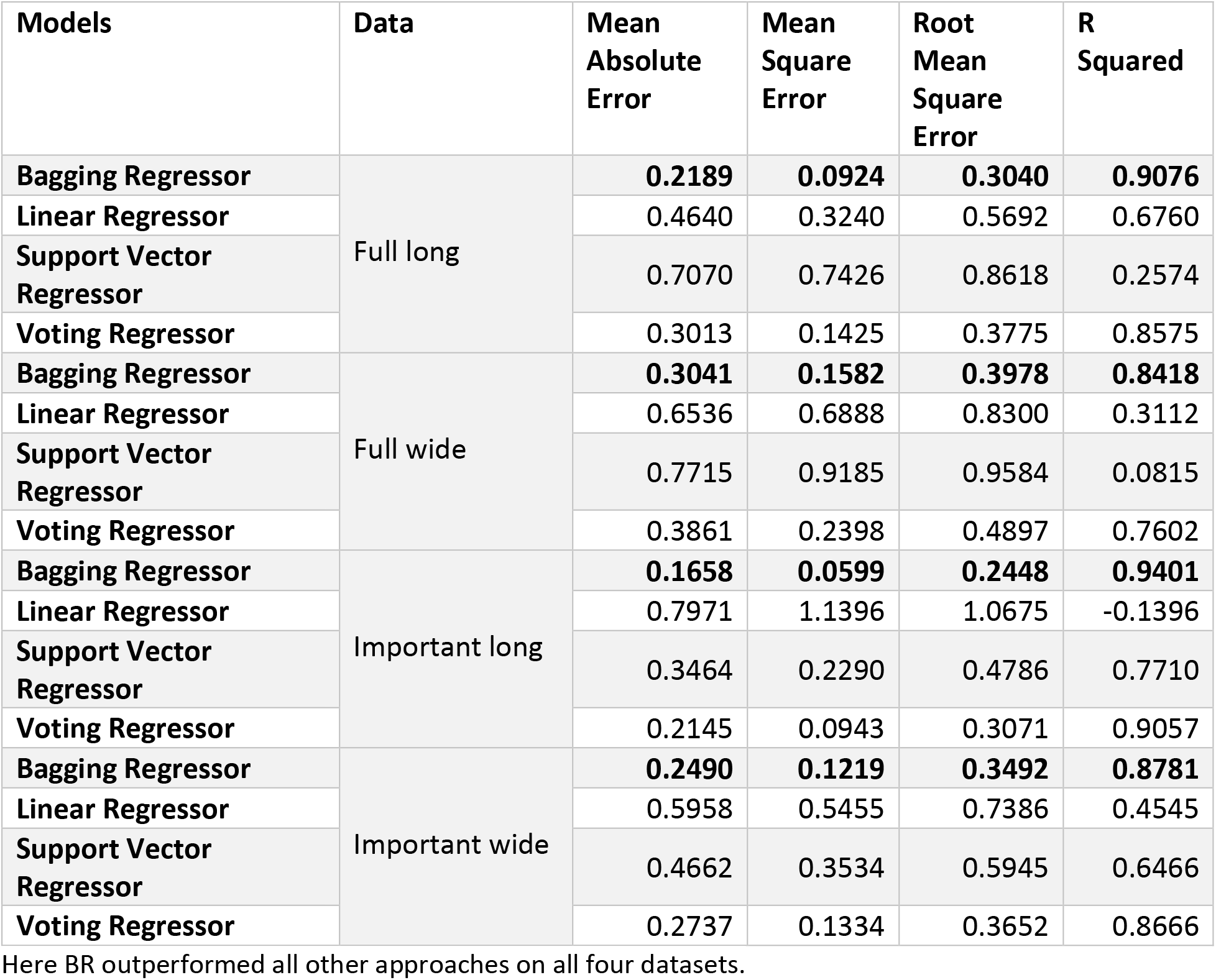
Base model accuracy for BR, LR, SVR, and VR approaches.

## Discussion

While previous research suggests that prior psychopathology, social adversity, and genomic variation at GRRN genes shape risk of traumatic stress, it remains unclear which of these factors figure most prominently in increasing prospective risk of PTSD. It is also clinically challenging to identify individuals at higher risk of PTSD because of its complex etiology and clinical course. Here we applied ML approaches to identify the features that associate with elevated prospective risk of PTSD. We used DNAm data from GRRN genes and multiple survey measures to determine the role of prior psychopathology, GRRN CpGs and social adversity as risk factors of future traumatic stress. ML identified prior PTSS as the most important predictor for prospective risk of PTSD. Many additional factors including PTS symptom clusters, loneliness, perceived discrimination and GRRN CpGs were identified as significant predictors for PTSD risk. Using these identified factors, ML models predicted the prospective risk of PTSD with high accuracy. We could not compare the accuracy of models in this study with existing approaches for PTSD (Dean et al., 2019; Karstoft et al., 2015; Schultebraucks et al., 2020; Wshah et al., 2019; Zhang et al., 2020) because this prior work used classification approaches, whereas we applied a regression task on a continuous outcome, PTSS, and different performance measures are used for classification and regression.

To apply our ML approach, we evaluated multiple models with different settings, i.e. base, and tuned models, on both long and wide datasets with different sets of features. We found that, in general, both base and tuned models perform very well, as confirmed using cross-validation on the training set and prediction on the final independent test set. Mean values on 10-fold cross-validation on the training sets showed the RF model to be the best performing model on the training set, followed by AB and GB. However, GB outperformed other methods using base model on test set. Results show that both GB and RF performed better, showing a lower error rate and higher R^2^ compared to other approaches, and the difference in prediction between the two is small. In the tuned model on test data, RF was the best model, but it was still less than the base model. Overall, all models showed better accuracy on the long vs. the wide dataset, both in terms of models applied using important and all features. One potential reason for this is the difference in sample size (210 in long vs. 148 in wide datasets). Another reason may be the missing values that are created when arranging the data in wide format. These missing values were imputed, but there is always some difference between the observed and imputed values. Two methods, SVR and VR, performed better using important features on the wide set as compared to the all the features on the long set but still less than the score on the important features on the long set. In general, SVR and LR performed poorly as compared to other approaches, whether used for long or wide data, full, or important features.

The results showed that not all features from the GRRN genes and phenotypes are needed to predict the risk of PTSS, or in other words, using only significant features increases the accuracy of prediction. The sets of 150 significant features produced better results when compared with other feature sets or all the features. It is evident that the significant sets of features both for the long and the wide datasets decreases the error rate and improves the prediction of the models. With this set, we identified significant CpGs and phenotypes as risk factors that predict the prospective risk of PTSS. From features identified as important in the long and wide datasets, we identified common features, where phenotypes were more numerous compared to CpGs.

From the list of significant features in phenotypes, PTSS in prior waves was highly predictive of prospective risk of PTSS. In particular, PTSS from wave 3 was of the highest importance in predicting wave 4 PTSS, confirming the importance of recency of symptom burden to prediction of future symptoms. This feature alone contributed >88 % of the significance in both long and wide approaches, and all 149 other features contributed to the remaining percentage. One interesting note is that the feature had almost the same score in both long and wide datasets. The symptom severity at wave 2 is also highly significant. These results are consistent with results of previous studies. For example, in a prospective study that followed adolescents and young adults for up to 50 months, 52% of the PTSD cases remitted during the follow-up period, whereas the remaining 48% showed no significant remission of PTSD (Perkonigg et al., 2005). The authors concluded that PTSD is usually a persistent and chronic disorder, and that symptom clusters might also be associated with the chronic course of PTSD (Perkonigg et al., 2005). Another study looking at PTSD, anxiety, and depression after the Spitak earthquake and political violence in Armenia showed no remission of PTSD over a 3-year interval, but that depression symptoms subsided over time (Goenjian et al., 2000). A longitudinal study showed that trauma-related psychopathology increases the risk of PTSD and significant impairment over time (Lewis et al., 2019). Our results also showed that prior post-traumatic psychopathology is highly important in predicting the prospective risk of PTSS in the subsequent year. We saw a decrease in the predictive significance of PTSS in relation to more distant symptoms, with wave 3 symptoms showing the highest importance in predicting wave 4 PTSS, followed by wave 2 and wave 1 symptoms. The higher importance of recent vs. more distant PTS symptoms in predicting future PTSS is consistent with a previous study that reported a general diminution in PTSD symptom severity over time (R. Yehuda et al., 2009). We also found prior depression and anxiety as significant predictors for PTSS risk, consistent with a prior ML study (Schultebraucks et al., 2020), which showed pre-deployment depression and anxiety as risk factors for PTSD in army personal deployed to Afghanistan. In addition, our results showed that all three PTSD symptom clusters (intrusion, avoidance, and hyperarousal), appear on the significant feature list. The symptom cluster, hyperarousal, that is detected as very significant in predicting PTSS by ML, has been associated and reported as the first symptoms to occur with chronic PTSD, followed by avoidance and intrusion symptoms (Bremner, Southwick, Darnell, & Charney, 1996). Our results suggest that symptom clusters contribute to the chronic course of PTSD and that people with higher symptom severities, both overall and across multiple symptom domains, are more likely to be at high risk of PTSD in the future.

In addition to these symptom-related findings, our ML results showed other phenotypes are significantly associated with predicting prospective risk of PTSD. For example, number of traumatic event types was identified as an important feature both overall and at waves 1 and 2, with similar levels of importance in both the long and wide datasets. More importantly, our analyses showed that the cumulative traumatic event type is a better predictor for PTSD risk as compared to traumatic event types from a single wave. Previous work has shown that cumulative traumatic burden is associated with increased risk of PTSD (Copeland, Keeler, Angold, & Costello, 2007; Ogle, Rubin, & Siegler, 2014), and additional work (Perkonigg et al., 2005) has reported that participants with a chronic course of PTSD are more likely to experience new traumatic events. Many additional social adversity factors, including loneliness, perceived discrimination, emotional mistreatment, were also predictive of prospective risk of PTSD in our analyses. Chronic PTSD has been associated with reduced social support, a higher frequency of social phobia, and greater avoidance symptoms (Davidson, Hughes, Blazer, & George, 1991). These results show that social adversity and trauma exposure contribute to the increased risk of PTSS even when biologic factors are taken into account, suggesting a multiplicity of mechanisms that explain the pathogenesis of PTSD.

Our work also identified GRRN CpG sites whose DNAm measures were predictive of prospective PTSD risk. There are 44 CpGs that were consistently significant in both long and wide analyses. Out of 44 CpGs, 3 CpGs (*cg04444450: NCOA2, cg20509117: IL6; LOC541472, cg05790989: POU2F1*) were significant in the long dataset and both the waves (wave 1 and wave 2) of the wide dataset. The significance indicates that the CpGs are highly important and consistent. All three of the genes related to these CpGs have been associated with PTSD. For example, Nuclear Receptor Coactivator 2 (*NCOA2*) has been identified as a biomarker for PTSD (Breen et al., 2019), and Interleukin 6 (*IL6*) has been associated with PTSD (Haxhibeqiri et al., 2019; Lima et al., 2019; Pervanidou et al., 2007; Somvanshi et al., 2020). POU Class 2 Homeobox 1 (*POU2F1*) has been implicated as a transcription factor located proximal to a SNP associated with emotional memory formation in PTSD (Wilker et al., 2018). *IL6* has been associated with depression and anxiety as well (Crawford et al., 2018; Ryan et al., 2017; Sanada et al., 2020). The CpGs *cg05616442* and *cg18071894* in gene AKT Serine/Threonine Kinase 1 (*AKT1*) showed highest importance score when looking at long and wide scores together. This gene has been associated with depression previously (Starnawska et al., 2019). The other CpGs on the significant list was *cg26495008* and *cg10913456*, both in the FKBP Prolyl Isomerase 5 (*FKBP5*) gene, previously associated with childhood maltreatment and depression (Klinger-König et al., 2019; Mehta et al., 2013); and risk of PTSD (Binder et al., 2008; Pape et al., 2018). The role of other significant genes detected by ML include: Nuclear Factor Kappa B Subunit 1 (*NFKB1*), which has been associated with personality disorders (Gescher et al., 2018); Interleukin 4 (*IL4*), which has been associated with PTSD (Smith et al., 2011); Nuclear Receptor Subfamily 3 Group C Member 1 (*NR3C1*), previously associated with emotion dysregulation, psychopathology (Cicchetti & Handley, 2017), depression (Borçoi et al., 2020; Efstathopoulos et al., 2018), and risk of PTSD (Schechter et al., 2015; Vukojevic et al., 2014); and Nuclear Factor Of Activated T Cells 1 (*NFATC1*), which has been previously associated with PTSD and depression (Kuan et al., 2017). It is clear from the literature that GRRN genes play a crucial role in mediating the stress response, and many genes identified here by our ML analyses have been previously associated with traumatic stress and stress-related psychopathology.

This study has some limitations: First, the sample size used in this study is limited, and a larger sample size may increase prediction accuracy and may help to design better generalized models. Second, many phenotype variables have missing data, a common issue in many domains. We imputed the missing values, but some differences may remain between imputed and the observed values. Missing values often represent hidden patterns in the data, and complete data may have provided more insights in this study. Future work involving a more complete dataset may address this issue. Second, the sample size used in this study is limited, and a larger sample size may increase prediction accuracy. Third, we could not assess the generalizability of the ML models due to the lack of an available independent, external dataset. In the future, the availability of such datasets will help to validate the ML models. Finally, in this dataset, we could not demonstrate the causal relationship of the identified predictors with the increased risk of PTSD, due to the limitations inherent to working human study participants.

Despite these limitations, this study has many strengths. The data used in this study is rich in terms of social adversity factors and biology, and there are very few large samples with the richness of data we have here and none, to our knowledge, that have leveraged such data for ML approaches in PTSD. In addition, we used standard pipeline and approaches for our analyses and implemented multiple ML models; these ML models are very efficient in predicting the prospective risk, confirmed by cross-validation, and using the independent test dataset. Our applied approach identified a specific list of DNAm features and phenotypes that play a significant role as predictors for prospective PTSD risk with high accuracy. These features chosen by ML as significant have been previously associated with PTSD. Our results suggest that ML can effectively use a wide variety of data such as DNAm, psychopathology, social adversity, and cell proportions to address the challenging task of identifying the associated risk factors and predicting PTSS risk. In conclusion, it is both challenging and crucial to identify people at higher risk of PTSD, and to understand the factors associated with elevating the prospective risk of traumatic stress. Our results show that ML approaches are efficient in identifying the factors that predict the prospective risk of PTSD with high accuracy. Many of the factors identified by ML as risk predictors are consistent with previous studies exploring the determinants of PTSD; our results extend this prior work, however, by assigning relative importance of these determinants from a wide range of social, psychopathological, and genomic features that has not been previously examined in the context of ML-based risk prediction of PTSD. Results from this study suggest that ML approaches may be further developed to detect elevated post-traumatic risk and efficiently to assist early intervention.

## Authors Contribution

**Agaz H Wani:** Conceptualization, Methodology, Analysis, Interpretation, Writing - original draft, review & editing. **Allison E. Aiello:** Conceptualization, Interpretation, Writing - review & editing. **Grace S. Kim**: Writing - review & editing. **Fei Xue**: Writing - review & editing. **Chantel L. Martin:** Writing - review & editing. **Andrew Ratanatharathorn:** Writing - review & editing. **Annie Qu:** Conceptualization, Writing - review & editing. **Karestan Koenen:** Conceptualization, Interpretation, Writing - review & editing. **Sandro Galea:** Conceptualization, Interpretation, Writing - review & editing. **Derek E. Wildman:** Conceptualization, Interpretation, Writing - review & editing. **Monica Uddin:** Conceptualization, Supervision, Interpretation, Writing - original draft, review & editing.

## Acknowledgments

We thank Mr. Zachary Graham and Miss. Sarah Burgan, who kindly helped in sample plating to generate data analyzed in this manuscript. We appreciate the time and effort of study participants, staff and volunteers of the Detroit Neighborhood Health Study.

