## Supplementary Material for "The Impact of Psychopathology, Social Adversity and Stress-relevant DNAm on Prospective Risk for Post-traumatic Stress: A Machine Learning Approach"

*Machine Learning Approaches*

Random Forest:

Random Forest (RF) is an ensemble learning approach that works by building several decision trees on random subsets of the data (Breiman, 2001). It then uses the average performance of each decision tree for the final prediction.

Adaboost:

Adaptive boosting, known as Adaboost (AB), is a technique used to build a highly accurate prediction model by using many models. In this approach, many models are built sequentially, each trying to correct its predecessor. More attention is paid to those training instances misclassified by the previous model (in order to correct them). This process results in adjusting misclassified cases sequentially multiple times, a process that ultimately produces a better model (Drucker, 1997; Freund & Schapire, 1997).

Gradient Boost:

Gradient Boost (GB) (Friedman, 2001) is another popular ensemble learning approach we used in this study. It works in a similar way to Adaboost, by building various sequential models. However, instead of adjusting the training instances on each step like AB, GB fits a new model on the residual error of the previous model, often resulting in a better prediction (less error).

Linear Regression:

Linear regression (LR) is one of the most straightforward approaches to model the relationship between independent and dependent variables. Linear regression fits a linear model to minimize the residual errors, i.e., the sum of squares between the observed target values and the predicted target values (Lai, Robbins, & Wei, 1978). In this study, we had multiple independent variables, so we used multiple linear regression.

Support Vector Regression:

The support vector machine (SVM) is a popular ML method used for classification problems. A variant of SVM, support vector regression (SVR) is used to deal with the regression problem, based on the principle of SVM. In this study, we used a linear, Gaussian Radial basis function (RBF) (Chang & Lin, 2011; Vapnik, 1995).

Bagging Regression:

The Bagging Regression (BR) approach trains the model on the random subsets of the data and then uses the prediction from each predictor to arrive at a final prediction (Breiman, 1996). We sampled the data with replacement (bootstrapping) and used soft voting (probability that an instance belongs to a certain class) for the final model prediction.

Voting Regression:

Voting regression (VR) is another ensemble learning approach that fits many models on the full dataset. The voting approach is then used for the individual prediction from each model to get the final prediction (An & Meng, 2010). In this study, as described above, we used six different models: RF, AB, GB, LR, SVR, and BR for the VR. In VR, a vote is obtained from each mentioned model, and a final prediction is made based on the maximum votes. Again, we used soft voting for the VR approach.

From the list of models mentioned above, we used RF, AB, and GB with two different settings: the model with default setting provided by the *Scikit-learn* framework (base model), and the model searching for best combinations of the parameters (tuned model) to verify if the prediction of the models improves.

*Evaluation measures:*

The evaluation metrics, mean absolute error (MAE), mean square error (MSE), root mean square error (RMSE), and R Squared ($R^2$) were used. MAE is the average absolute difference between the observed and the predicted values in the data. It gives the measure of the magnitude of the error, without considering the direction. MSE is similar to MAE except that it is the average of the squares of error between the actual and the predicted values. MSE is always non-negative, and value close to zero means a better prediction. RMSE is the measure of the square root of the prediction error. Lower RMSE indicates a better fit of the model. Finally, $R^2$ is the proportion of the variance explained by the independent variables for the dependent variable. It is defined as a percentage and has a value between zero and one, zero meaning no fit (0%), and one meaning perfect fit (100%).

*Analysis*:

As we have DNAm data from two-time points (waves) for some participants, we performed two types of analyseis. First, we arranged DNAm data row-wise; this produced a matrix with more than one row of DNAm data for participants with more than one time point of DNAm data. Second, we arranged DNAm data column-wise; this produced a matrix with a single row for each participant and multiple columns of DNAm for a particular CpG site, for the participants with more than one time point of DNAm data. Arranging data in a row-wise format increases the sample size and may thus help in better prediction. Analysis using the column-wise data may help find the DNAm signatures important in both time-points to predict the risk of PTSS. In both analyses we used wave specific phenotype data, including prior psychopathology such as PTSS, anxiety and depression symptom severity, and PTSS symptom clusters.

| #Features | Mean Absolute Error | Root Mean Square Error |
|---:|---:|---:|
| **10** | 0.2080 | 0.3032 |
| **30** | 0.2021 | 0.2896 |
| **50** | 0.1952 | 0.2745 |
| **70** | 0.2037 | 0.2902 |
| **90** | 0.2004 | 0.2910 |
| **110** | 0.2015 | 0.2947 |
| **130** | 0.2084 | 0.2994 |
| **150** | 0.1939 | 0.2850 |
| **170** | 0.2106 | 0.3035 |
| **190** | 0.2098 | 0.3033 |
| **210** | 0.2145 | 0.3070 |
| **230** | 0.2060 | 0.3032 |
| **250** | 0.2188 | 0.3148 |
| **270** | 0.2182 | 0.3088 |
| **290** | 0.2146 | 0.3038 |
| **310** | 0.2199 | 0.3105 |
| **330** | 0.2189 | 0.3107 |
| **350** | 0.2241 | 0.3135 |
| **370** | 0.2225 | 0.3151 |
| **390** | 0.2280 | 0.3174 |
| **410** | 0.2309 | 0.3227 |
| **430** | 0.2322 | 0.3200 |
| **450** | 0.2316 | 0.3224 |
| **470** | 0.2271 | 0.3178 |
| **490** | 0.2228 | 0.3164 |
| **510** | 0.2296 | 0.3208 |
| **530** | 0.2292 | 0.3220 |
| **550** | 0.2330 | 0.3239 |
| **570** | 0.2288 | 0.3194 |
| **590** | 0.2353 | 0.3277 |

**Table S1:** Different sets of features with mean absolute error (MAE) and root mean square error (RMSE). It shows the mean values of error rates of RF, AB, and GB. The set of 150 features shows better accuracy.

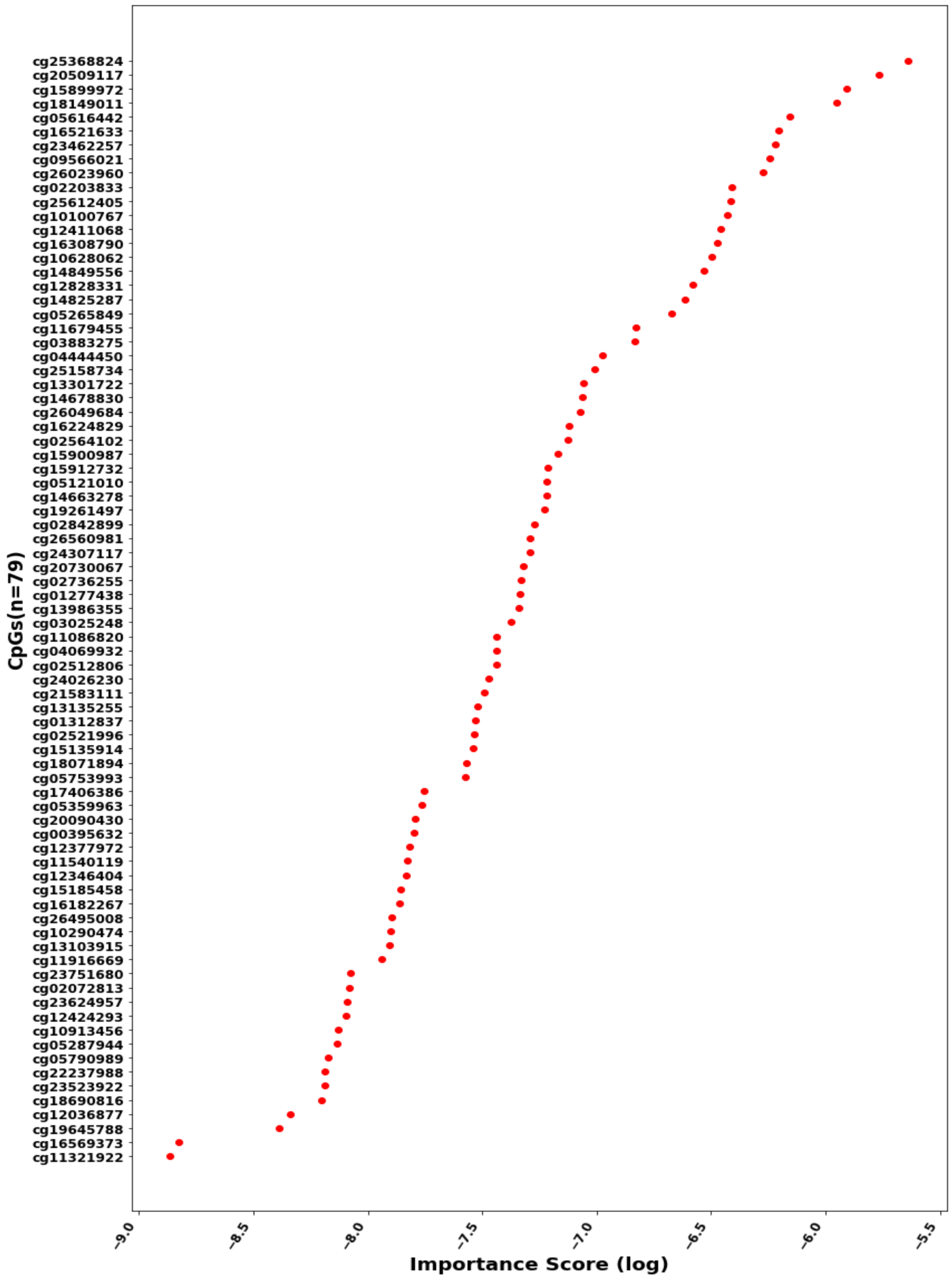


**Figure S1:** Significant CpGs and importance score in log scale from the long data.

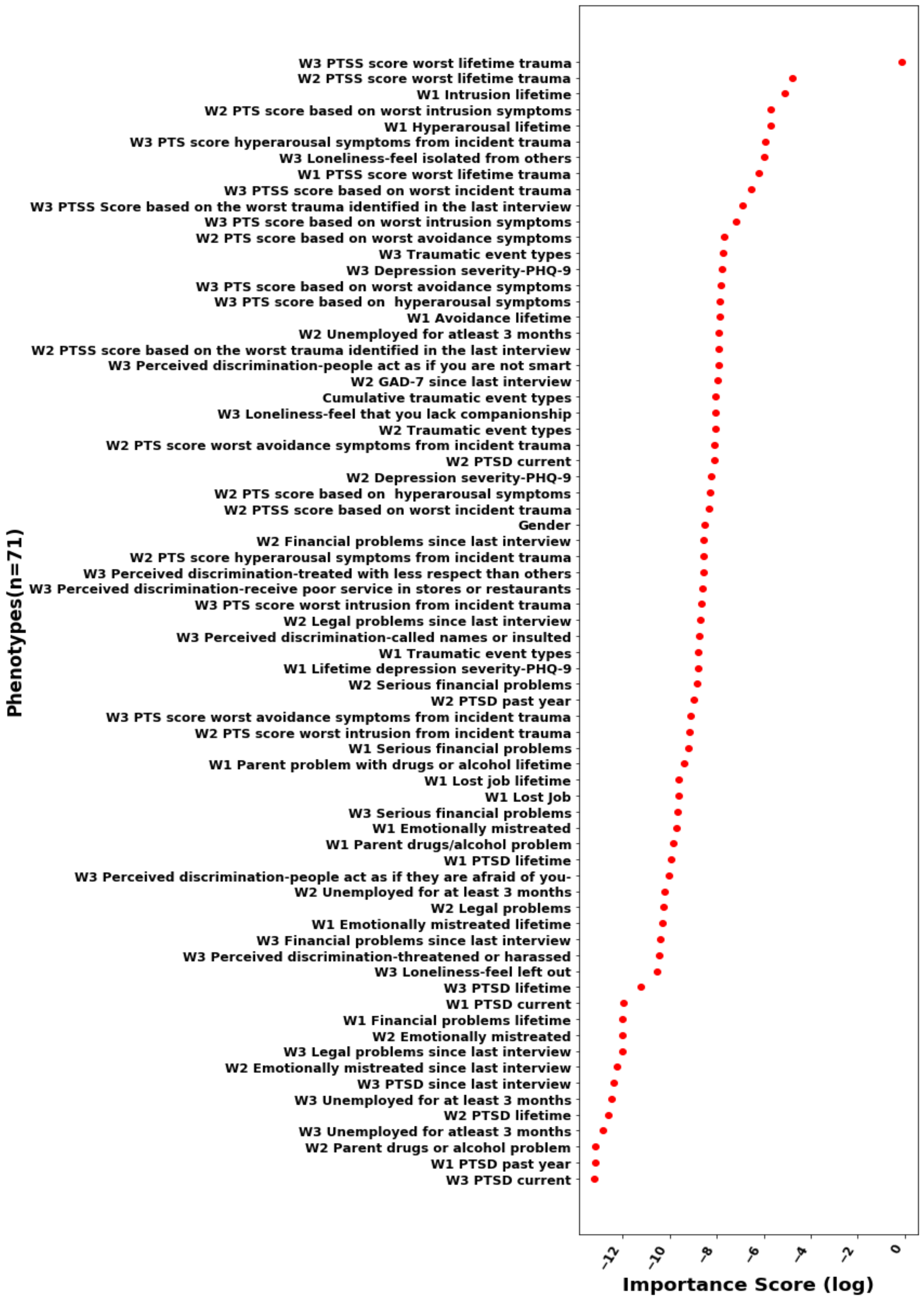


**Figure S2:** Significant phenotypes and importance score in log scale from the long data.

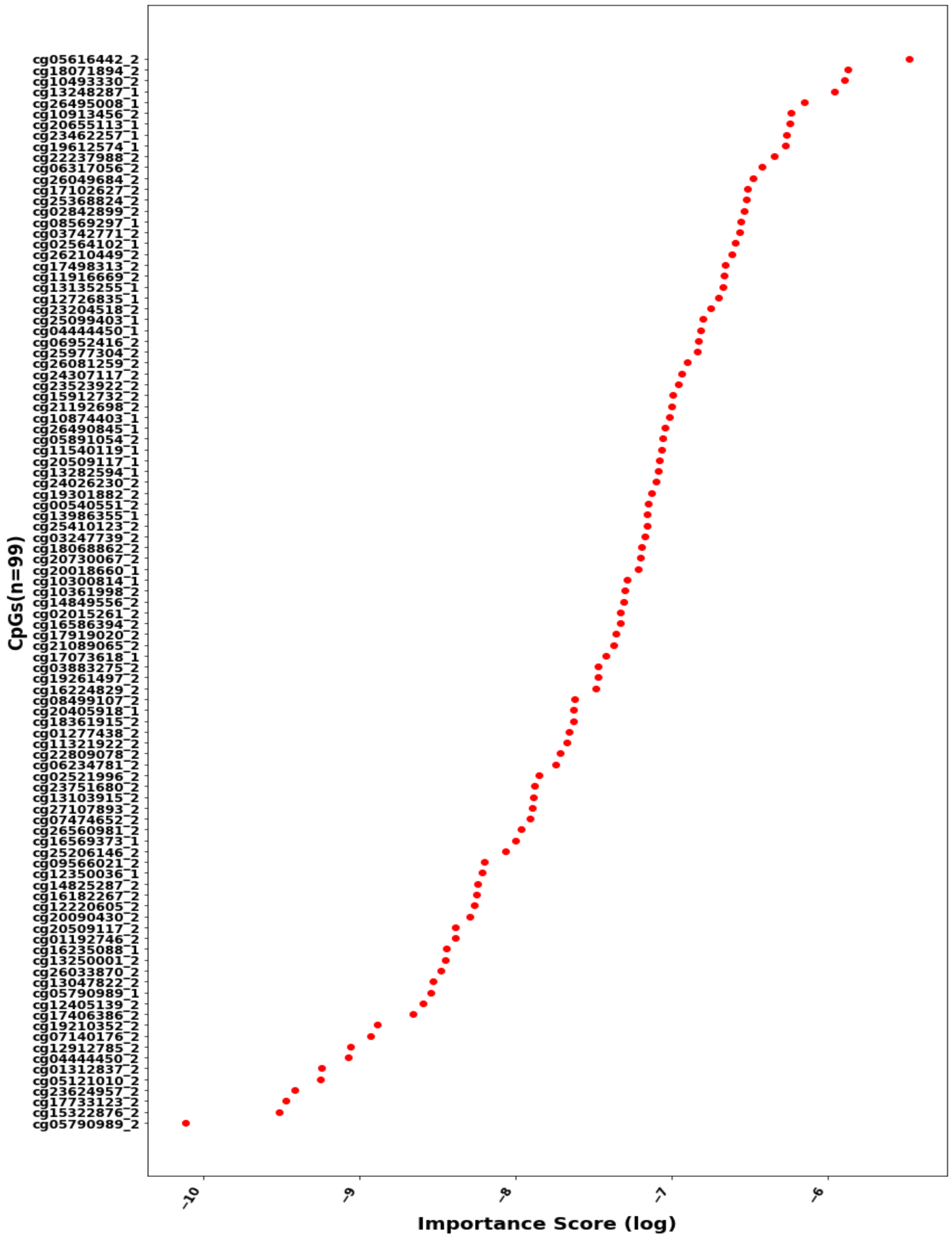


**Figure S3:** Significant CpGs and importance score in log scale on the wide data. The set of 99 CpGs has representation from both waves. The CpGs on Y-axis with '_1' are from wave 1, and '_2' are from wave 2.

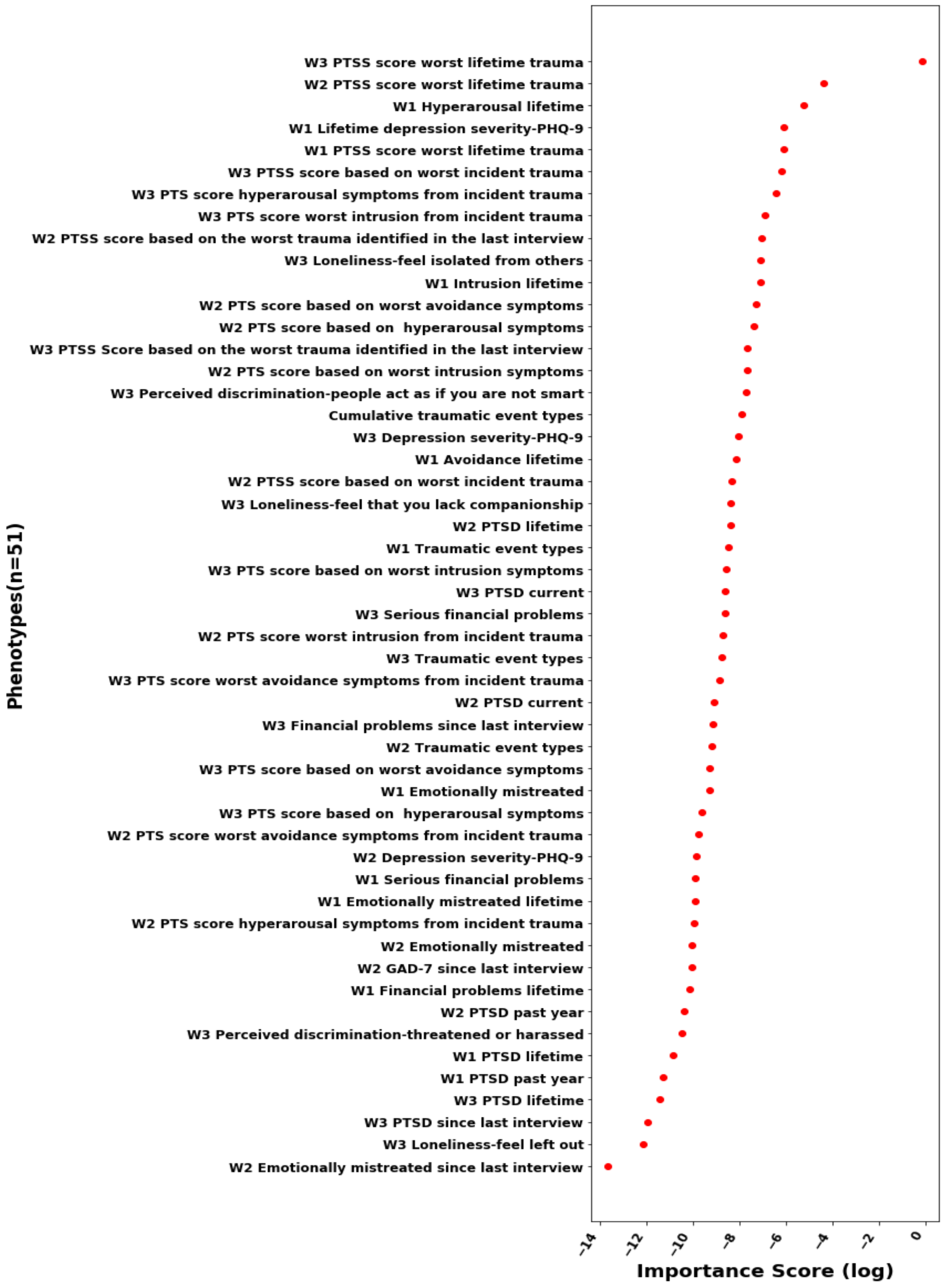


**Figure S4:** Significant phenotypes and importance from the wide data format.

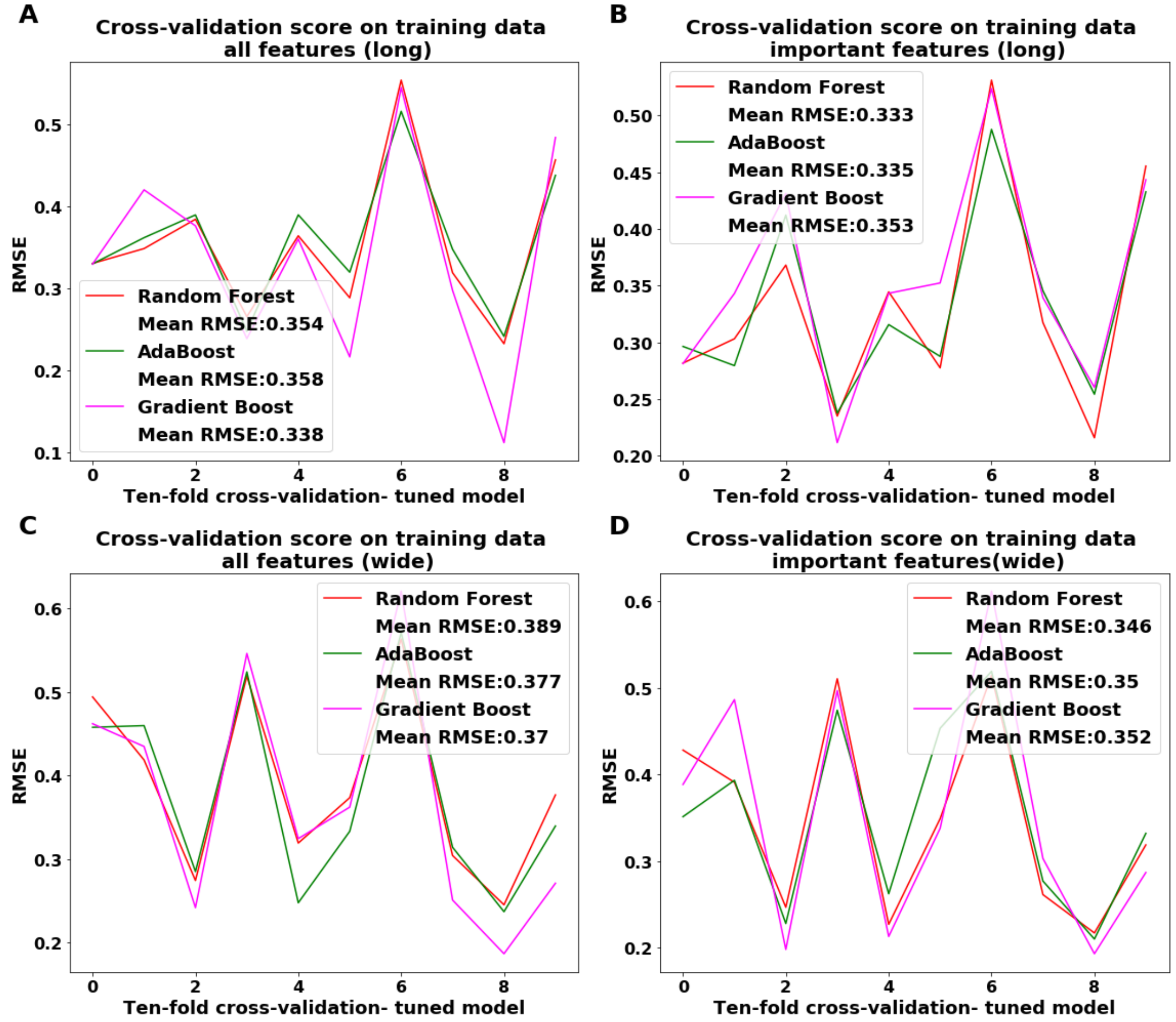


**Figure S5:** RMSE scores of the tuned models

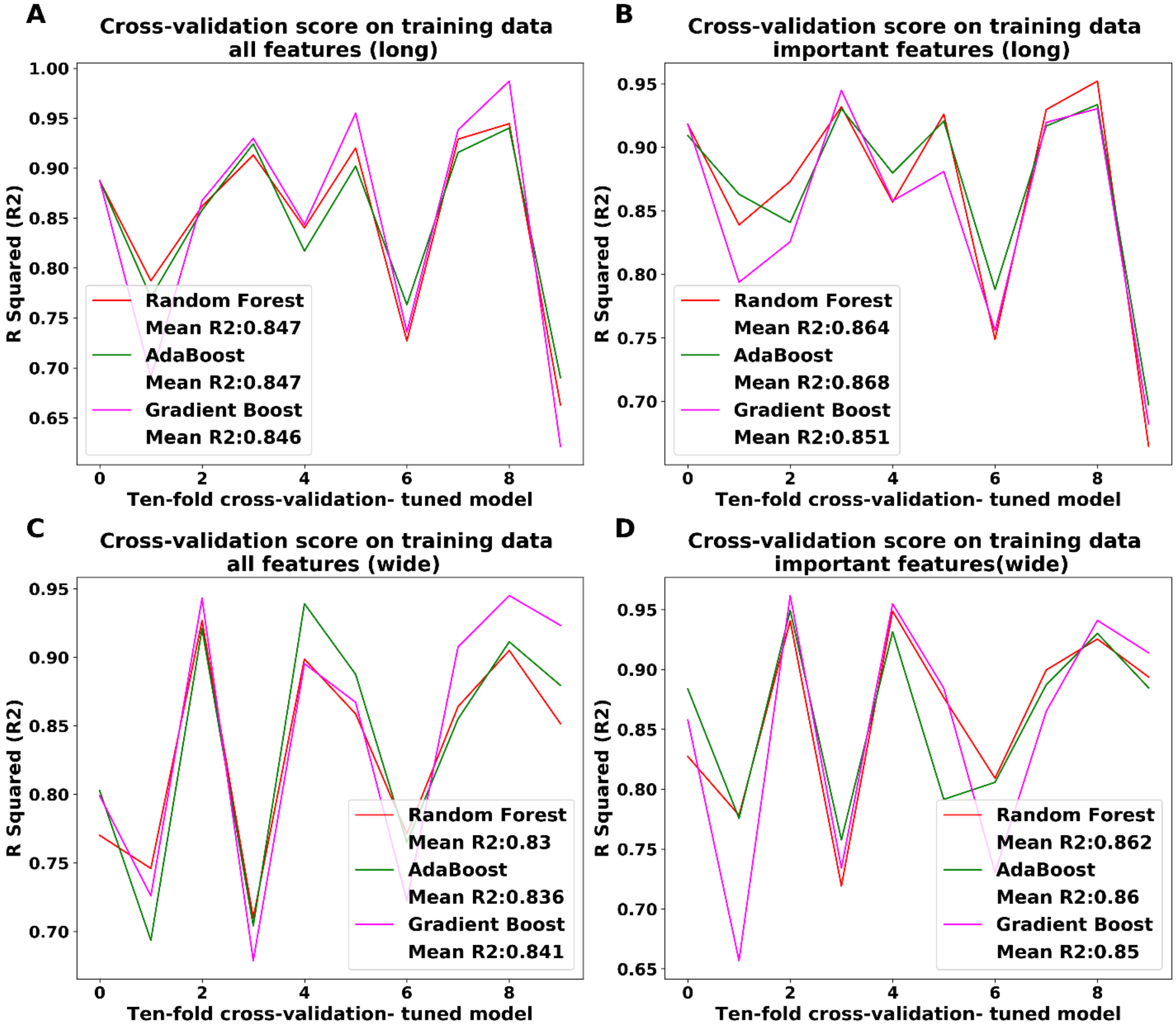


**Figure S6:** $R^2$ score of the tuned models
